# A highly responsive pyruvate sensor reveals pathway-regulatory role of the mitochondrial pyruvate carrier MPC

**DOI:** 10.1101/611806

**Authors:** R Arce-Molina, F Cortés-Molina, PY Sandoval, A Galaz, K Alegría, S Schirmeier, LF Barros, A San Martín

## Abstract

Mitochondria generate ATP and building blocks for cell growth and regeneration, using pyruvate as the main substrate. Here we introduce PyronicSF, a user-friendly GFP-based sensor of improved dynamic range that enables real-time subcellular quantitation of mitochondrial pyruvate transport, concentration and flux. We report that cultured mouse astrocytes maintain mitochondrial pyruvate in the low micromolar range, below cytosolic pyruvate, which means that the mitochondrial pyruvate carrier MPC controls the decision between respiration and anaplerosis/gluconeogenesis in an ultrasensitive fashion. The functionality of the sensor in living tissue is demonstrated in the brain of *Drosophila melanogaster* larvae. Mitochondrial subpopulations are known to coexist within a given cell, which differ in their morphology, mobility, membrane potential, and vicinity to other organelles. The present tool can be used to investigate how mitochondrial diversity relates to metabolism, to study the role of MPC in disease, and to screen for small-molecule MPC modulators.

## Introduction

Mitochondria are the chief energy generators of animal cells, accounting for over 90% of ATP production, and they also generate building blocks for the synthesis of sugars, amino acids, nucleic acids and prosthetic groups, essential elements for tissue growth, plasticity and regeneration. In addition to their metabolic functions, mitochondria are involved in diverse physiological and pathophysiological processes, including Ca^2+^ signaling, the production of reactive oxygen species, aging and degeneration, cell death and oncogenesis. Some open issues in mitochondrial physiology are the regulation of intermediate metabolism, the coordination between cytosolic and mitochondrial pathways, the decision between catabolism and anabolism, and the crosstalk between mitochondria and other organelles like plasma membrane and endoplasmic reticulum. Transversal to these questions is the meaning of mitochondrial diversity.

A major mitochondrial substrate for mammalian cells is pyruvate, a 3-carbon organic acid produced from glucose, lactate and amino acids. Pyruvate enters mitochondria through the mitochondrial pyruvate carrier (MPC) ^1–3^. Once in the matrix, the flux of pyruvate is split. Carbon is either shed to generate ATP via oxidative phosphorylation or alternatively, carbon is accrued to generate oxaloacetate, a process termed anaplerosis, which provides building blocks for biosynthesis and constitutes the first step of gluconeogenesis.

Pyronic, the first genetically-encoded sensor for pyruvate ^4^, has permitted the measurement of cytosolic pyruvate in several organisms with high temporal resolution, see for example ^5–11^. This article introduces a new version of Pyronic that improves on the original sensor in terms of dynamic range and facility of use, as it is imaged with the 488 nm Argon laser of standard confocal microscopes. To demonstrate the usefulness of the probe, we measured cytosolic, nuclear and mitochondrial pyruvate concentration, MPC-mediated permeability and determined the metabolic flux of small groups of mitochondria. The main finding of this study is that mitochondrial pyruvate lies in the low micromolar range, endowing the MPC with ultrasensitive control over anaplerosis. Experiments were also carried out in fruit fly larvae to demonstrate pyruvate dynamics in living tissue.

## RESULTS

### A highly responsive pyruvate sensor

Figure 1A illustrates PyronicSF (Single Fluorophore), in which the bacterial transcription factor PdhR ^12^ has been linked to a circularly-permuted version of GFP (cpGFP) ^13^ (DNA sequence in Supplementary Fig. S1). Exposure of the sensor to pyruvate *in vitro* caused a strong increase in fluorescence when excited by blue light (Fig. 1B). Excited at 488 nm, the increase in fluorescence emission was ≈ 250%, with a K_D_ of 480 μM (Fig. 1C). As expected from the behavior of PdhR ^4^ in the FRET sensor Pyronic, PyronicSF was insensitive to lactate, to other structurally related organic acids, and to NAD^+^ and NADH (Fig. 1D). While the sensing of pyruvate by PyronicSF was insensitive to pH, there was a pH-dependent shift in the whole dose-response curve (Supplementary Fig. S2). Because the FRET sensor Pyronic is insensitive to pH ^4^, we attribute the effect of pH on PyronicSF to the pH-sensitivity typical of cpGFPs ^14, 15^. When required, the signal can be corrected by using parallel measurements with a pH probe or by taking advantage of the pH-sensitivity of PyronicSF excited at its isosbestic point (435 nm; Fig. 1B), as detailed in Supplementary Fig. S2. Excited with violet light, pyruvate caused a decrease in fluorescence emission (Fig. 1B), which may be exploited to obtain ratiometric measurements using, for example, the 405 nm laser line available in some confocal microscopes. Expressed in the cytosol of HEK293 cells, PyronicSF showed a homogeneous distribution and responded to a saturating pyruvate load with a change in fluorescence similar to that of the purified protein (Fig. 1E). The amplitude of the response of PyronicSF was > 6 times that of Pyronic (Fig. 1E) ^4^.

**Figure 1.**
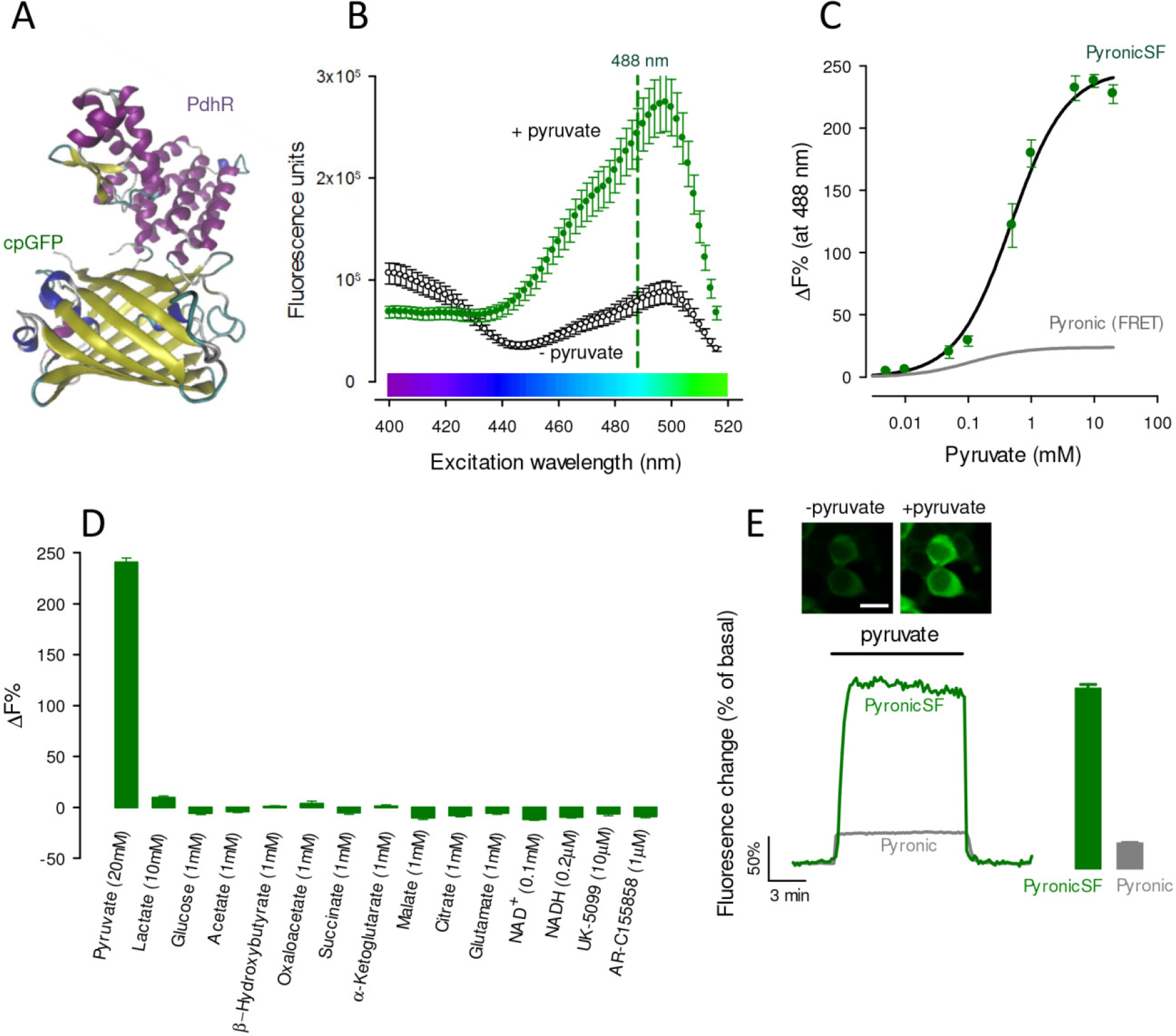
Characterization of PyronicSF. (A) PyronicSF. cpGFP flanked by linkers was inserted between aminoacid residues 188 and 189 of PdhR. DNA sequence in Supplementary Fig. S1). (B) Excitation spectra of PyronicSF in the absence and presence of 10 mM pyruvate. Data are mean ± s.e.m. from 3 protein extracts. (C) PyronicSF emission (488 nm excitation) as a function of pyruvate concentration. Data are mean ± s.e.m. from 3 protein extracts. The best fit of a rectangular hyperbola to the data is shown, K_D_ = 480 ± 65 μM, maximum fluorescence change was 247%. The *in vitro* saturation curve of the FRET sensor Pyronic is plotted in gray ^4^. (D) PyronicSF emission in the presence of metabolites and transport inhibitors. Data are mean ± s.e.m. from 3 protein extracts. (E) Pyruvate dynamics in mammalian cells. HEK293 cells expressing PyronicSF were exposed to 10 mM pyruvate. Images show cells before and during exposure to pyruvate. Bar represents 20 μm. Bar graphs summarize data (mean ± s.e.m.) from 54 cells in four experiments (PyronicSF), and 59 cells in five experiments (Pyronic).

### Estimation of MPC activity

PyronicSF was targeted to the mitochondrial matrix using the destination sequence of cytochrome oxidase. The targeting was successful in various cell types, as evidenced by the labeling of elongated cytoplasmic structures that are of shape and size characteristic of mitochondria (Supplementary Fig. S3). To test the functionality of the sensor we chose astrocytes, cells that combine oxidative phosphorylation with anaplerosis, and that in culture are very thin, ideal to resolve mitochondria (Fig. 2A). Confirmation of correct targeting was provided by colocalization with the voltage-sensitive dye TMRM (Supplementary Fig. S4). Mito-PyronicSF also colocalized with the red fluorescent protein mCherry targeted to mitochondria (Fig. 2A and Supplementary Fig. S3). Exposure of astrocytes co-expressing mito-PyronicSF and mito-mCherry to pyruvate resulted in a reversible increase in fluorescence ratio, evidencing mitochondrial uptake (Fig. 2B). The rise in mitochondria was much slower than its accumulation in the cytosol, monitored indirectly with PyronicSF targeted to the nucleus (Supplementary Fig. S5). In cells in which mito-PyronicSF was expressed by lipid transfection (as opposed to adenoviral transduction), there was sometimes a diffuse staining explained by inefficient mitochondrial targeting/retention. In such “leaky” cells, the increase in signal elicited by a pyruvate load was biphasic, with a rapid phase attributable to the cytosolic sensor, followed by a slow mitochondrial phase (Supplementary Fig. S5). The FRET sensor Pyronic could also be targeted to mitochondria in functional form (Supplementary Figs. S3 and S6), but four copies of the destination sequence were needed, probably because of its larger size relative to PyronicSF.

**Figure 2.**
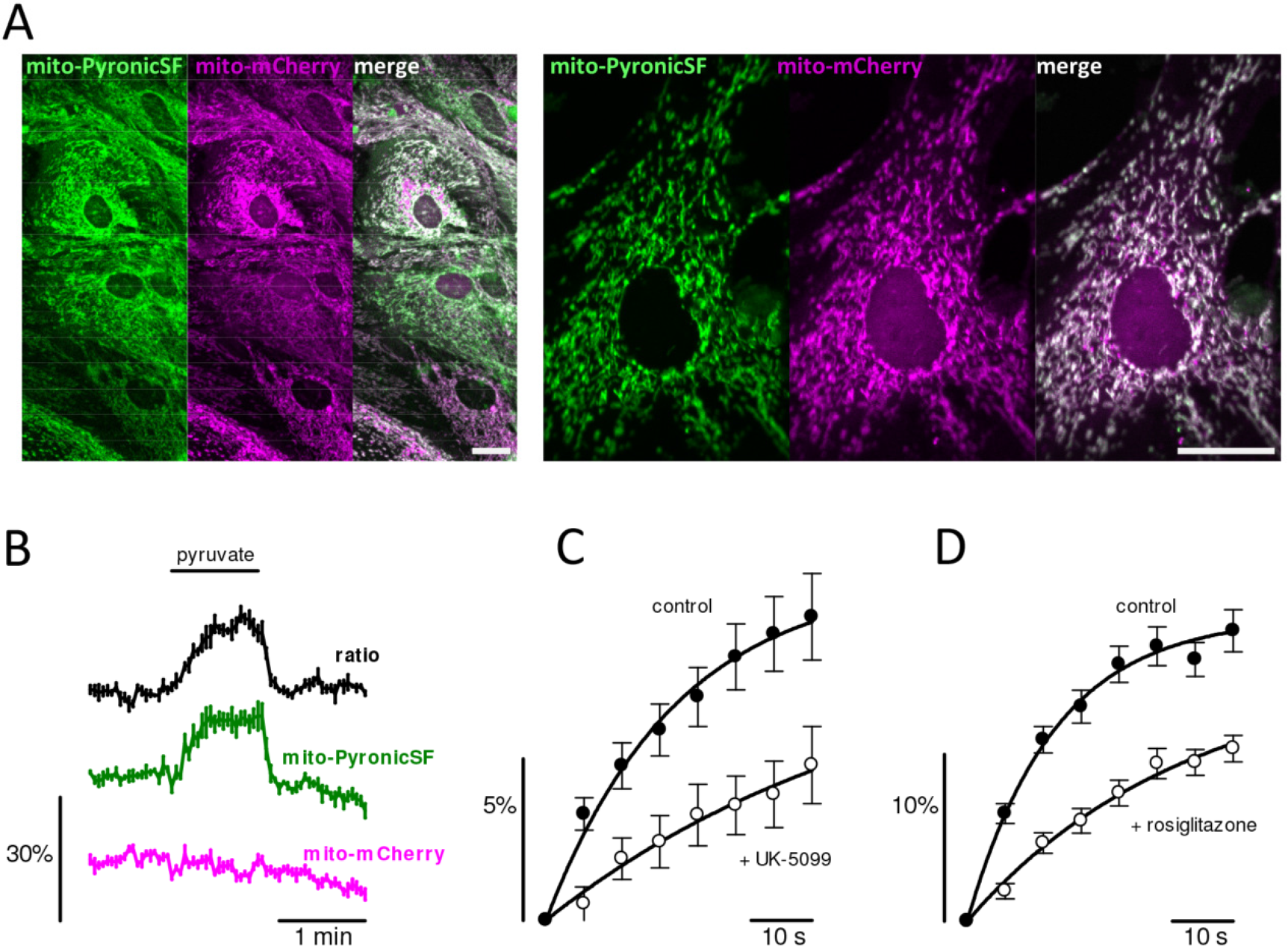
MPC-mediated mitochondrial pyruvate transport in astrocytes. (A) Astrocytes co-expressing mito-PyronicSF (green) and mito-mCherry (magenta). Bars represents 10 μm. (B) Cultures were exposed to 3 mM pyruvate. Data corresponds to mean ± s.e.m. (4 cells in a representative experiment). (C) Cultures were exposed to 3 mM pyruvate in the absence (black symbols) and presence of 10 μM UK-5099 (white symbols). Data are mean ± s.e.m. of 31 cells from eight experiments. Initial rates (%/min), estimated by fitting a single exponential function to the data (continuous lines), were 32 ± 4 (control) and 10 ± 5 (UK-5099). (D) Cultures were exposed to 3 mM pyruvate in the absence (black symbols) and presence of 30 μM rosiglitazone (white symbols). Data are mean ± s.e.m. of 51 cells from nine experiments. Initial rates (%/min), estimated by fitting a single exponential function to the data (continuous lines), were 78 ± 6 (control) and 26 ± 3 (rosiglitazone).

The uptake of pyruvate by mitochondria was sensitive to the specific MPC1 blocker UK-5099 ^1–3, 16^(Fig. 2C). The average degree of uptake inhibition was 69 % (Fig. 2C). Possible explanations for the incomplete inhibition may be variations in the subunit composition of the MPC ^16^ and/or the presence of alternative routes. This type of uptake protocol seems amenable to the identification and characterization of pharmacological inhibitors of the MPC, a transporter that has been singled out as a target of clinical interest ^6, 17–23^. To illustrate the potential of mito-PyronicSF for drug discovery, we show here that the insulin sensitizer rosiglitazone inhibited the uptake of pyruvate by astrocytic mitochondria by 67% (Fig. 2D), providing direct confirmation that MPC is a target of thiazolidinediones ^17^.

### Quantification of mitochondrial pyruvate concentration

The concentration of pyruvate in the mitochondrial matrix of intact cells is unknown. Given the 1:1 stoichiometry between proton and pyruvate transport by the MPC, and the respective pH gradient between cytosol and mitochondria (7.2 and 7.8 ^24^, equivalent to a 4-fold proton gradient), mitochondrial pyruvate may reside anywhere between a few micromolar and 4 times the cytosolic concentration of pyruvate, something that is also not known with precision. At extracellular levels of glucose, lactate and pyruvate found in brain tissue, steady-state mitochondrial pyruvate was found to lie at the low end of the detection range of PyronicSF (Figs. 3A and 3C), which is in agreement with parallel measurements using the FRET sensor (Figs. 3B and 3C). A one-point calibration of both sensors was accomplished by forcing a nominal zero cytosolic pyruvate by MCT-accelerated exchange with lactate ^4, 25^, followed by interpolation in the dose-response curve obtained *in vitro* (Fig. 1C). Using this protocol, median mitochondrial pyruvate measured with PyronicSF was 21 μM. Experiments with the sensor targeted to the cytosol showed that the median steady-state concentration of pyruvate in this compartment was 33 μM. Estimated with the FRET sensor, median pyruvate concentrations were 25 μM in mitochondria and 40 μM in cytosol. Thus, both sensors revealed that mitochondrial pyruvate is lower than cytosolic pyruvate.

**Figure 3.**
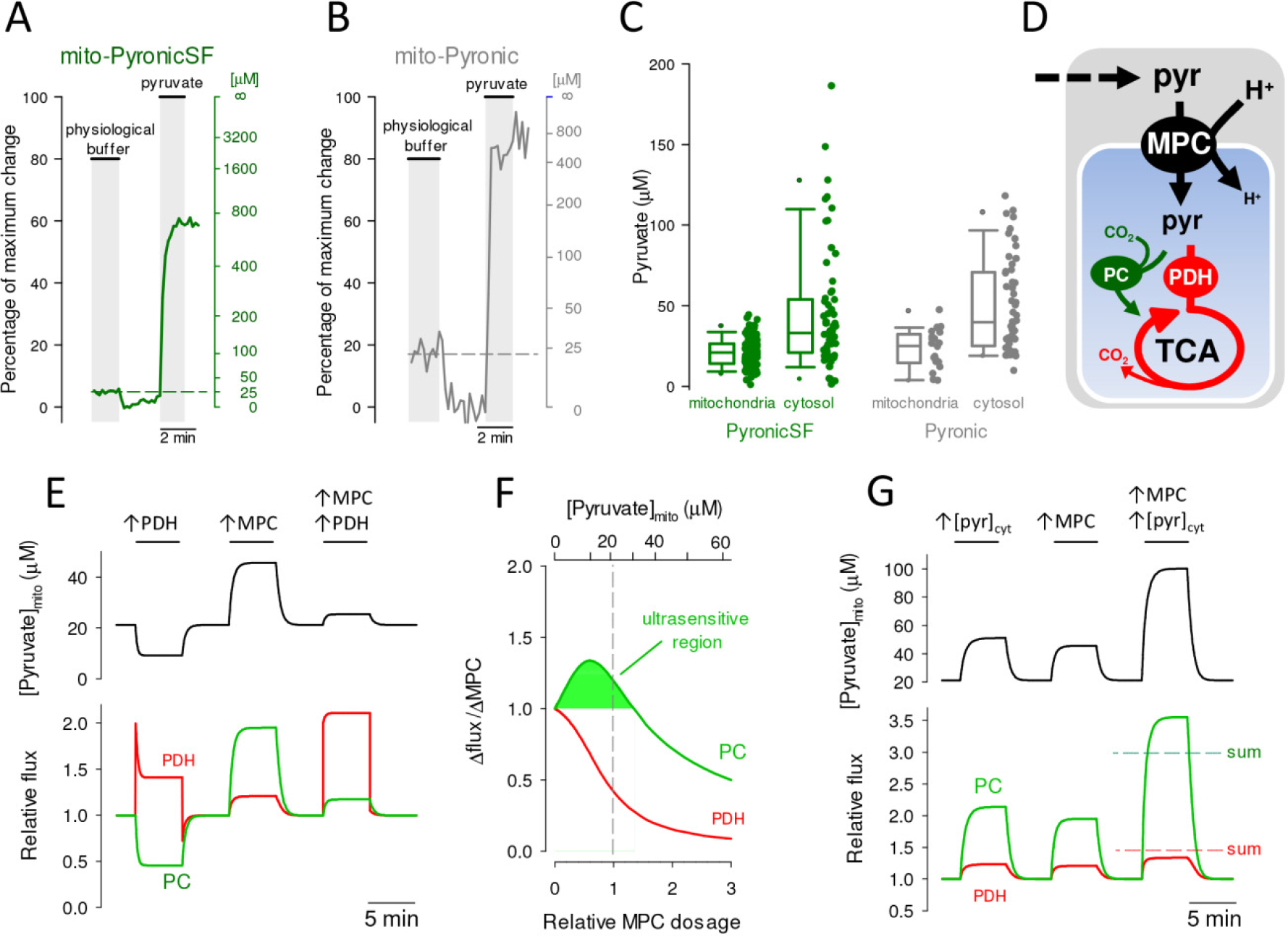
Steady-state mitochondrial and cytosolic pyruvate. (A) Astrocytes expressing PyronicSF or Pyronic in mitochondria or cytosol were first incubated in a buffer containing physiological concentrations of glucose (2 mM), lactate (2 mM) and pyruvate (0.2 mM), followed by removal of pyruvate by accelerated-exchange with 10 mM lactate and exposure to 10 mM pyruvate. Representative trace from a single astrocyte expressing mito-PyronicSF. Data are shown as percentage of the maximum change (left) and pyruvate concentration (right), with reference to the response of the sensor obtained *in vitro* (Fig. 1C). (B) Representative trace from a single astrocyte expressing mito-Pyronic. Data are shown as percentage of the maximum change (left) and pyruvate concentration (right), with reference to the response of the sensor obtained *in vitro* ^4^. (C) Steady-state mitochondrial and cytosolic pyruvate concentrations measured with PyronicSF or Pyronic at physiological concentrations of glucose, lactate and pyruvate, as illustrated in panels A and B. Data are from 131 cells in ten experiments (mito-PyronicSF), 59 cells in four experiments (cytosolic PyronicSF), 20 cells in seven experiments (mito-Pyronic), and 50 cells in five experiments (cytosolic Pyronic). (D) Mitochondrial pyruvate dynamics. Pyruvate enters mitochondria (blue compartment) and is metabolized by PDH and the tricarboxylic acid cycle (TCA), or is carboxylated by PC. (E) Simulation of pyruvate dynamics in response to PDH and MPC modulation. The effects of activating PDH and MPC by 100% on mitochondrial pyruvate concentration (top panel) and on the fluxes of PDH and PC (bottom panel) are shown. Steady-state cytosolic and mitochondrial pyruvate were 33 μM and 21 μM. Cytosolic and mitochondrial pH were 7.2 and 7.8. Steady-state PDH and PC fluxes were 0.91 and 0.29 μM/s. (F) Ultrasensitive modulation of PC flux by the MPC. The curves show the degree of flux increase at PC and PDH relative to the degree of MPC activation. The shaded area under the PC curve indicates the range of pyruvate concentrations at which PC flux increases more than 1% when MPC is activated by 1%. MPC dosage was normalized at 3.24 μM. (G) Synergic effect of cytosolic pyruvate and MPC activity on PC flux. The effects of increasing cytosolic pyruvate and MPC activity by 100% on mitochondrial pyruvate concentration (top panel) and on the fluxes of PDH and PC (bottom panel) are shown. The sums of the independent effects are indicated by interrupted lines.

### Astrocytic mitochondrial pyruvate is at optimal level for MPC control of anaplerosis

With knowledge of pyruvate levels inside and outside mitochondria, a mathematical model could be built to appraise the role of the MPC on catabolism and anabolism. The two main mitochondrial pyruvate sinks are pyruvate dehydrogenase (PDH; EC 1.2.4.1), which is in charge of pyruvate catabolism towards energy production, and pyruvate carboxylase (PC; EC 6.4.1.1), which catalyzes pyruvate anabolism towards anaplerosis and synthesis (Fig. 3D). Their combined flux was set at 1.2 μM/s ^26^, with PDH and PC respectively accounting for 76% and 24% ^27^. Next, the dosage of MPC was tuned so that at a cytosolic pyruvate of 33 μM and the pH gradient estimated previously in these cells ^24^, mitochondrial pyruvate stabilized at 21 μM, as determined above. With the system configured in this way a number of observations could be made. Firstly, activation of the MPC is necessary for efficient stimulation of oxidative metabolism. For example, PDH activation does not translate into a sustained flux increase because mitochondrial pyruvate becomes depleted, leading to partial PDH desaturation (Fig. 3E). More dramatically, the fall of pyruvate leads to a strong decrease of flux through PC, a “steal” flux phenomenon. However, if the activation of PDH is accompanied by activation of the MPC, the flux through PDH can be sustained, with minimal disturbance of PC flux. Figure 3E also shows that MPC modulation is sufficient to efficiently modulate the flux through PC. Actually, Fig. 3F shows that if the steady-state concentration of mitochondrial pyruvate is below 30 μM, a given change in MPC activity will be amplified in terms of PC flux, a phenomenon that has been termed ultrasensitivity ^28^. For example, at 21 μM mitochondrial pyruvate, a 1% MPC increase causes a 1.2% increase in PC flux. In contrast, PDH flux increases by only 0.4% (Fig. 3F). A similar analysis showed that PC flux is also highly responsive to cytosolic pyruvate. For example a rise in cytosolic pyruvate, as may result from glycolytic activation, is faithfully followed by an increase in PC flux and a minor increase in PDH flux (Fig. 3G). Remarkably, the MPC and cytosolic pyruvate interact synergistically with respect to PC flux but antagonistically with respect to PDH flux. A coincidental rise of MPC activity and cytosolic pyruvate will cause an increase in PC flux larger than the sum of the two independent effects, but for PDH the effects are not even additive (Fig. 3G). The exquisite sensitivity of PC flux to MPC activity (and to cytosolic pyruvate) is not observed at high mitochondrial pyruvate levels (Fig. 3F).

### Quantification of mitochondrial pyruvate flux

A protocol to approach flux was devised based on the idea that acute inhibition of mitochondrial pyruvate entry at the MPC should cause a progressive fall in mitochondrial pyruvate as it is consumed by PDH and PC. Analogous methods have been applied successfully to the measurement of whole cell glucose, lactate and pyruvate consumptions in various cell types ^9, 29, 30^. As shown in Fig. 4A, MPC stoppage with UK-5099 caused immediate depletion of mitochondrial pyruvate. Consistent with the partial UK-5099 inhibition of the pyruvate transport described above, the depletion was not complete. We do not think that the partial effect is explained by insufficient UK-5099, because similar steady-states were reached at 0.5 μM and 10 μM UK-5099 (53 ± 12% versus 47 ± 6%; 12 cells in three paired experiments, p > 0.05, data not shown) and the half-inhibition constant for MPC inhibition by UK-5099 is only 50 nM ^31^. Thus, the present protocol underestimates actual the rate of mitochondrial pyruvate consumption by an approximate factor of 2. Reportedly, long term exposure to UK-5099 causes irreversible inhibition of the MPC ^17, 32^ but this was not observed at the low concentrations and short exposure times used here (Fig. 4B). This reversibility opens the possibility of before-and-after experiments. Using such a protocol, uncoupling of oxidative phosphorylation (OXPHOS) strongly stimulated stimulated mitochondrial pyruvate consumption (Fig. 4C) whereas conversely, inhibition of OXPHOS with the cytochrome oxidase inhibitor azide resulted in a lower rate (Fig. 4D). A representative example of the estimation of mitochondrial pyruvate consumption expressed in absolute terms is shown in Fig. 4E. There was substantial variability between cells, which ranged from 0.29 to 3.1 μM/s, with a median value of 1.1 μM/s (Fig. 4E).

**Figure 4.**
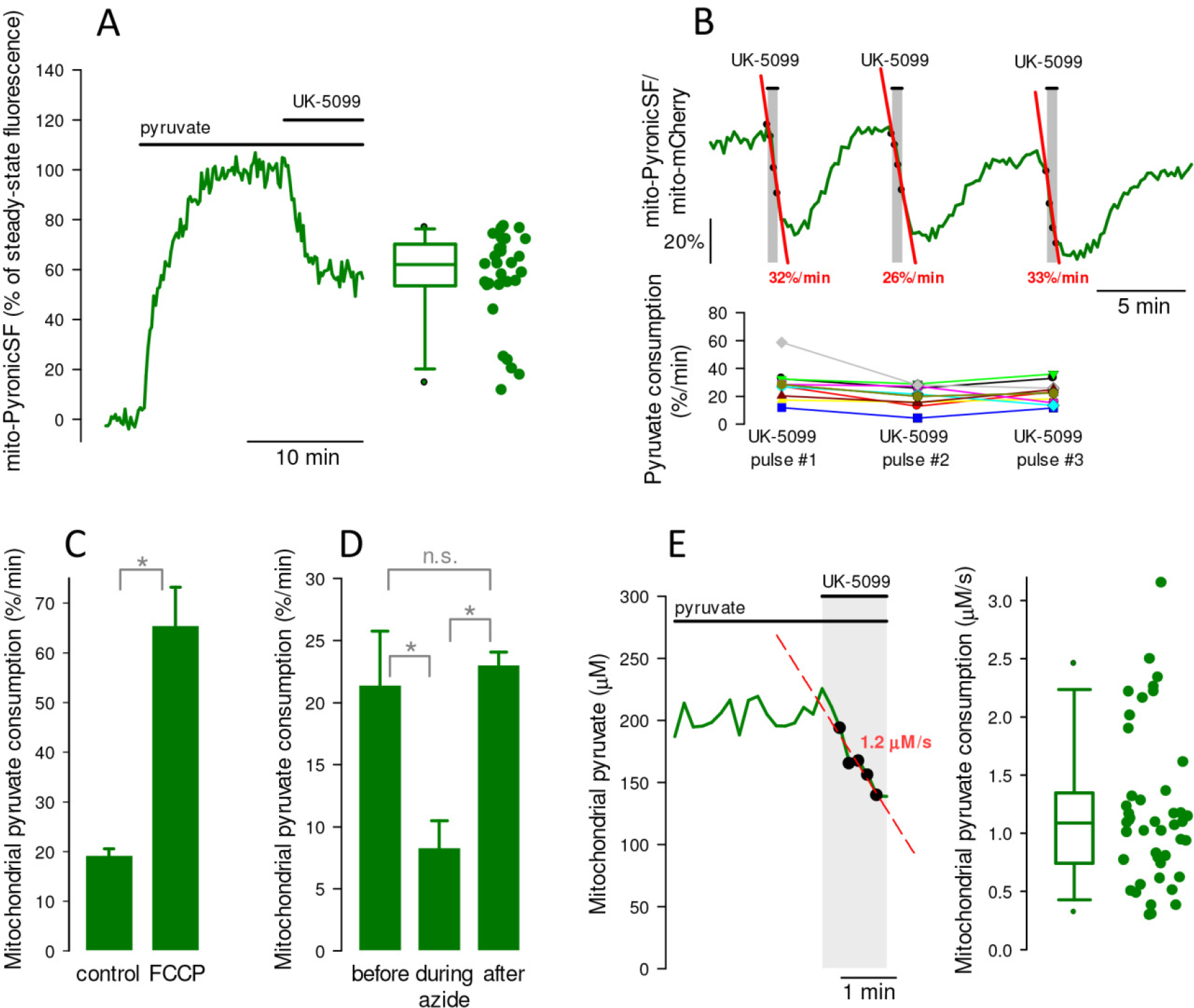
Measurement of mitochondrial pyruvate consumption rate in individual cells. (A) Time course of mitochondrial pyruvate level after an astrocyte was exposed 5 mM pyruvate and then to 10 μM UK-5099. The new steady-state is represented on the right, as percentage of the level before addition of UK-5099. Median = 62%. Data are from 29 cells in ten experiments. (B) An astrocyte incubated in 5 mM pyruvate was exposed three times for 30 s to 0.5 μM UK-5099. Rates of pyruvate depletion are shown in red. The result of three similar experiments (9 cells) is shown below. (C) The rate of mitochondrial pyruvate depletion induced with 10 μM UK-5099 was monitored before and after exposure to the proton ionophore FCCP (1 μM). Data are mean ± s.e.m (29 cells in three experiments). (D) The rate of mitochondrial pyruvate depletion induced with 10 μM UK-5099 was monitored before, during and after exposure to the cytochrome oxidase inhibitor azide (5 mM; 16 cells in three experiments). (E) An astrocyte superfused with 3 mM pyruvate was exposed to 10 μM UK-5099, resulting in a rate of depletion of 1.2 μM/s. The right panel represents the summary of sixteen experiments (44 cells).

To exemplify how mito-PyronicSF might serve to study subcellular metabolism, pyruvate concentration and consumption were determined in mitochondria located in different regions of a cell. Steady-state pyruvate in a small group of peripheral mitochondria was found to be higher than in mitochondria lying close to the nucleus. Application of the transporter-block protocol showed that pyruvate consumption may also differ within a cell (Fig. 5).

**Figure 5.**
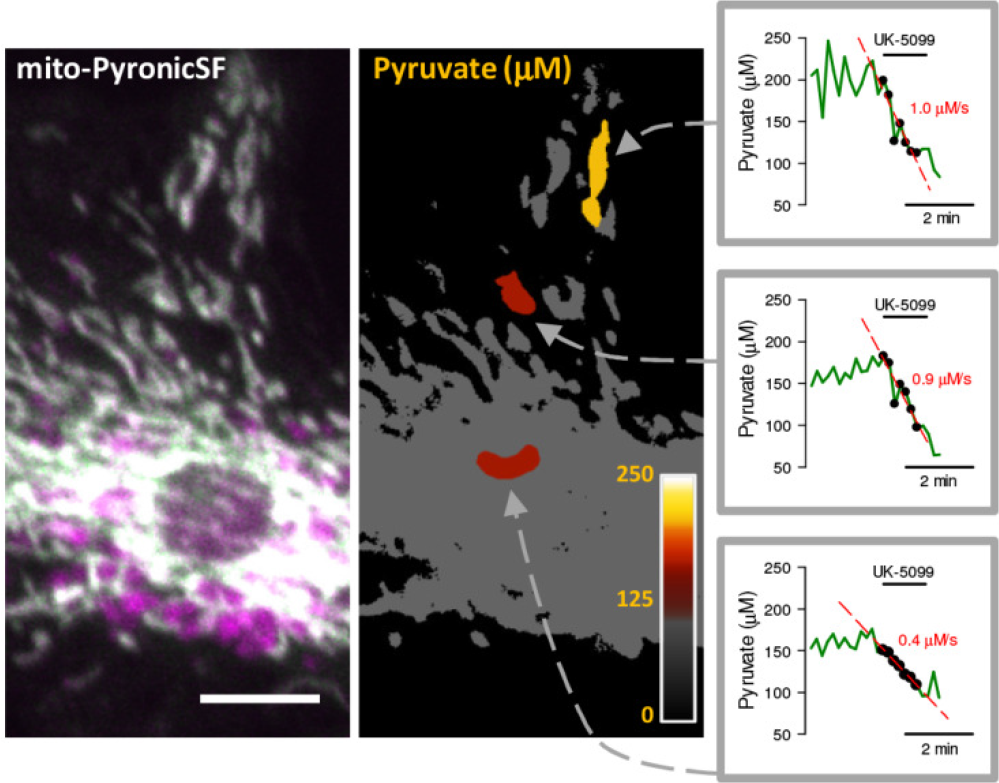
Pyruvate concentration and consumption in discrete mitochondria. Mito-PyronicSF (gray) in an astrocyte expressing mito-PyronicSF and mito-mCherry (left). Bar represents 10 μm. The righthand image shows three regions of interest colored according to the look-up table in the inset (0 to 250 μM pyruvate, calibrated as described for Fig. 3). The three graphs show pyruvate consumption rates in the regions of interest, determined with 10 μM UK-5099 in cells incubated with 3 mM pyruvate.

### Mitochondrial pyruvate dynamics in brain tissue

The ability of mito-PyronicSF to monitor pyruvate in living tissue was investigated in *Drosophila melanogaster* larvae. For facility of access, we studied perineurial glial cells, which form a monolayer separating the brain from the surrounding hemolymph. PyronicSF expressed very well in cytosol and mitochondria of these cells (Figs. 6A-B). Superfusion of acutely isolated brains with pyruvate resulted in a quick increase in cytosolic pyruvate, revealing the presence of abundant surface pyruvate transporters in these cells (Fig. 6C-E). The response of mitochondria was slower and plateaued at lower pyruvate levels, consistent with mitochondria being a site of pyruvate consumption downstream of the cytosol (Figs. 6D-E). In the presence of a buffer containing glucose, lactate and pyruvate, the steady-state level of pyruvate was much higher in the cytosol than in mitochondria (Fig. 6C-D). Experiments are planned to measure transmitochondrial pyruvate and pH gradients in the presence of normal hemolymph substrates. Nevertheless, the steep transmitochondrial pyruvate gradient measured here suggests that the MPC is also a key regulator of the balance between catabolism and anabolism in perineurial glial cells.

**Figure 6.**
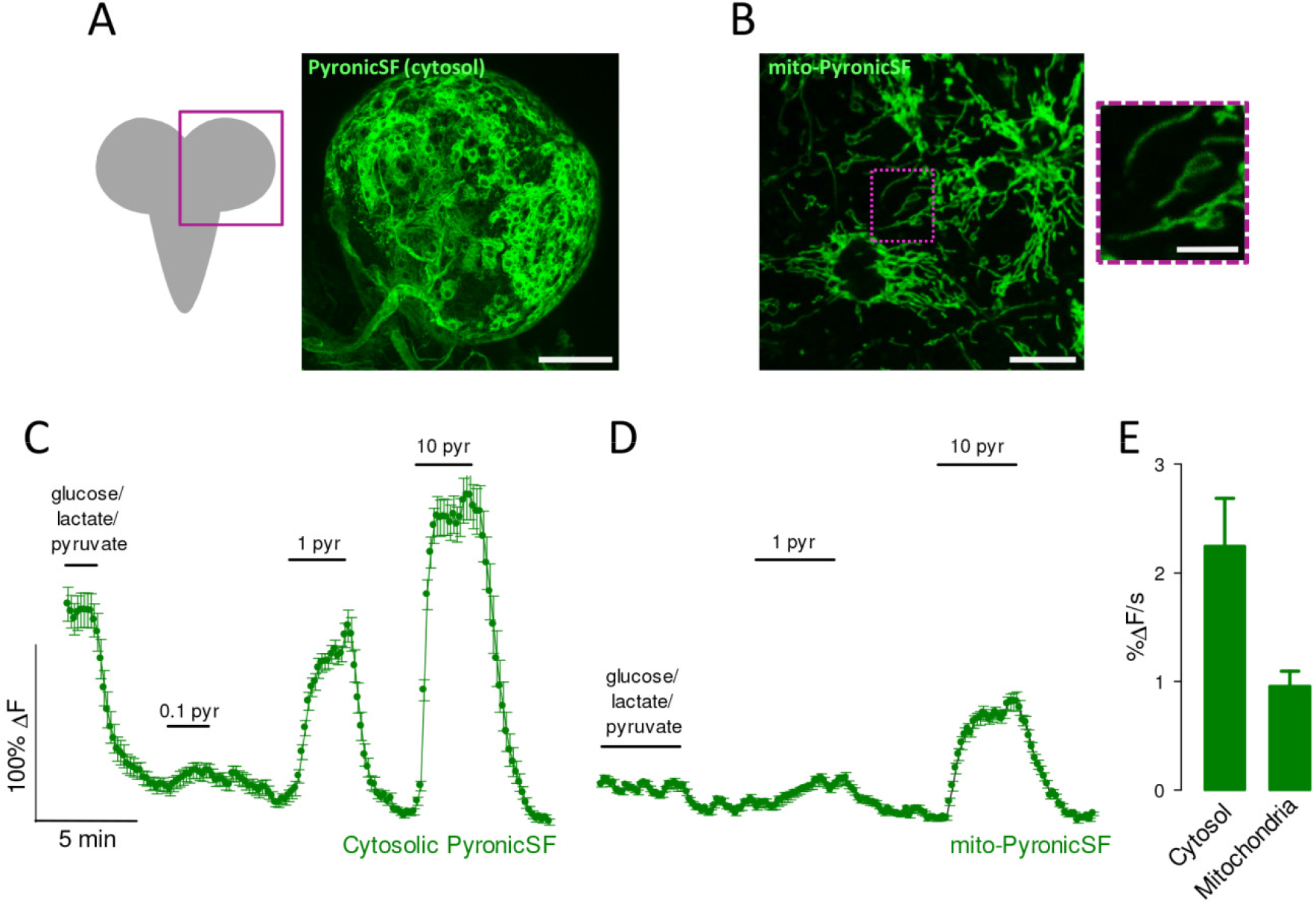
Pyruvate dynamics in glial cells of *Drosophila melanogaster*. Brains were acutely dissected from *Drosophila melanogaster* larvae expressing PyronicSF in the cytosol or mitochondria of perineurial glial cells. (A) PyronicSF in the cytosol of perineurial cells. Bar represents 100 μm. (B) Mito-PyronicSF in perineurial cells. Bar represents 10 μm. An area containing clearly identifiable mitochondria is shown under higher magnification on the right. Bar represents 5 μm. (C) A brain expressing cytosolic PyronicSF in perineurial cells was superfused with HL3 buffer containing 5 mM glucose, 1 mM lactate and 0.5 mM pyruvate. After removal of the substrates, the tissue was sequentially exposed to 0.1, 1 and 10 mM pyruvate. Data are mean ± s.e.m. (20 cells). (D) A brain expressing mito-PyronicSF in perineurial cells was superfused with HL3 buffer containing 5 mM glucose, 1 mM lactate and 0.5 mM pyruvate. After removal of the substrates, the tissue was sequentially exposed to 1 and 10 mM pyruvate. Data are mean ± s.e.m. (20 cells). (E) Rates of PyronicSF fluorescence increase in response to 10 mM pyruvate. Data are mean ± s.e.m. (60 cells from three experiments similar to those shown in C-D).

## DISCUSSION

The present article introduces PyronicSF, a new tool for the study of metabolism. PyronicSF is genetically-encoded and compatible with standard fluorescence microscopes. In combination with *ad-hoc* protocols, this sensor permits the measurement of transport, concentration and flux of pyruvate in intact mitochondria. In combination with suitable experimental models, PyronicSF may be adapted to the analysis of intact organs, cell populations, single cells or even individual mitochondria. Demonstrating its potential, we showed that in mouse astrocytes and probably in perineurial cells from *Drosophila melanogaster*, mitochondrial pyruvate lies in the low micromolar range, a finding that informs about the regulatory role of the MPC.

### A new approach to mitochondrial function

Mitochondria are the chief ATP producers of eukaryotic cells and also a principal site for the generation of building blocks for macromolecular synthesis. The speeds of these catabolic and anabolic pathways are assessed by diverse techniques ^30^ of complementary strengths and weaknesses. For catabolism, respirometry is the standard technique since the days of Otto Warburg. The rate of oxygen consumption, which is tightly linked to the production of ATP, is specific and quantitative but requires cell populations, typically more than 10,000 cells in modern devices. Isotopic techniques were introduced later on to map specific metabolic pathways, although with low temporal resolution. They also require many cells. These direct measures of metabolic flux have been complemented by fluorescent indicators of pH, membrane potential, free radicals and calcium. Thanks to the spatiotemporal resolution of fluorescence microscopy they afford measurements in the range of seconds and the ability to resolve up to single organelles, but are not informative about the speed of metabolism ^33^. A few years ago, our group introduced the FRET sensor Pyronic ^4^ that has permitted high resolution measurement of pyruvate in multiple contexts from cultured cells to living animals ^5–11^. The present sensor PyronicSF improves on the original sensor on several grounds: i. It has a higher dynamic range. ii. It does not require FRET detection technology, so it can be used in standard wide-field and confocal microscopes. iii. It is smaller and therefore more easily targetable to subcellular domains. iv. Because of its lower affinity for pyruvate, when expressed in mitochondria it provides an extended window for rate monitoring. v. It can estimate mitochondrial pyruvate consumption under physiological conditions (i.e. in the presence of glucose). vi. It permits monitoring pyruvate concentration, permeability and flux in individual mitochondria, thus offering a practical tool for the functional study of mitochondrial heterogeneity. One limitation of PyronicSF is its pH sensitivity, which can be corrected as explained in the Results Section. In principle, buffering by a sensor may interfere with fast ligand dynamics, as happens with Ca^2+ 34^, but this problem is not anticipated for pyruvate because its levels are higher (μM instead of nM) and metabolic transients are slow (seconds instead of ms). Possible interference of surface permeability in the study of MPC transport activity may be circumvented using permeabilized cells. Assays based on mito-Pyronic may be combined with the study of the MPC using luminescence ^6^.

### Ultrasensitive regulation of anaplerosis/gluconeogenesis by the MPC

Why is the mitochondrial concentration of pyruvate important? After it became established that pyruvate enters mitochondria through a transporter and not via simple diffusion ^35, 36^, issue was raised as to the role of the transporter on pyruvate utilization ^37, 38^. This is not a trivial question, as some transporters play important roles in flux control, for example glutamate transporters in the brain, and others do not, such as monocarboxylate carriers in most tissues ^39^. Key to this controlling role is the thermodynamic gradient driving the transporter. For the MPC, the relevant parameters are pH and pyruvate. It is well established that there is steep transmitochondrial proton gradient, which ranges between 0.6 and 1 pH units ^24, 40^, but for pyruvate, the numbers have not been available. Pyruvate in samples of brain tissue obtained by quick-freezing was measured at 99 μM ^41^, although fast glycogen degradation during the procedure may have affected the result ^42, 43^. The concentrations found here in cultured astrocytes showed that in these cells the MPC is poised to modulate metabolic decisions. Would that be the case in other cell types? Gluconeogenesis from lactate and amino acids in liver feeds the brain with glucose during fasting and exercise, but becomes excessive in diabetes, leading to hyperglycemia and chronic illness. Gluconeogenesis starts with the carboxylation of pyruvate by pyruvate carboxylase (PC) and this step may have a more important role in rate modulation than anticipated ^20, 44^. The MPC has also been pinpointed as a target in neurodegenerative diseases, because its inhibition mobilizes glutamate towards oxidation therefore ameliorating excitotoxicity ^22^. In contrast, in cancer cells, MPC acts as a protective factor by inhibiting the Warburg effect ^18^ and MPC inhibition enhanced tumor aggression and resistance to chemo- and radiotherapy ^18, 23^. Growth under MPC-inhibition is sustained by reprogramming of mitochondrial metabolism including enhanced glutaminolysis and lipid catabolism ^19^. Knowledge of mitochondrial pyruvate concentration, permeability and flux a cellular and subcellular levels may contribute to the understanding of these complex diseases, as well as in the identification and characterization of drugs that target the MPC, such as thiazoledinediones ^17^.

## ACKNOWLEDGEMENTS

We thank Karen Everett for critical reading of the manuscript. This work was partly supported by Fondecyt grants 11150930 to ASM and 1160317 to LFB, and Deutsche Forschungsgemeinshaft (DFG) grants SFB1009 and SCHI 1380/2 to SS. The Centro de Estudios Científicos (CECs) is funded by the Chilean Government through the Centers of Excellence Basal Financing Program of CONICYT.

## AUTHOR CONTRIBUTIONS

R.A-M. designed and performed most of the experiments in cultured cells, analyzed data, prepared figures, contributed to mathematical modeling and drafted the text. F.C-M. performed experiments *in vitro* and in cultured cells. P.Y.S. and K.A. contributed to experiments in cultured cells. A.G. performed experiments *in vitro*. S.S. generated transgenic lines and contributed to experiments in *Drosophila*. L.F.B. conceived and designed experiments *in vitro* and in cultured cells, contributed to mathematical modeling and drafted the text. A.S.M designed the sensor, planned and performed experiments and helped with drafting the manuscript. All authors discussed the data and critically revised the manuscript.

## METHODS

Standard reagents were acquired from Sigma and Merck.

### Generation and *in vitro* characterization of PyronicSF

Single fluorophore pyruvate sensors were built using the bacterial transcription factor PdhR ^12^ and cpGFP ^13^. To obtain purified protein, the DNA-coding sequence was cloned into pGST-Paralell1 ^45^ and transformed into competent E. coli BL21 (DE3). A single colony was grown in LB medium with 100 mg/ml ampicillin and expression was induced with 400 μM IPTG. Cells were collected by centrifugation at 5000 rpm (4 °C) for 10 min and disrupted by sonication (Hielscher Ultrasound Technology) in 5 mL of Tris-HCl buffer pH 8.0. A cell-free extract was obtained by centrifugation at 10,000 rpm (4 °C) for 1 hour. Proteins were purified using a glutathione resin as recommended by the manufacturer (GST Buffer and Resin Kit, General Electric Healthcare Life Sciences). After cleavage of GST with protease TEV (AustralProteins), eluted proteins were incubated overnight at 4 °C and measured with a microplate reader analyzer (EnVision, PerkinElmer). Excitation spectra were obtained collecting emission at 515 ± 15 nm. The variant that showed the largest change in fluorescence intensity upon binding pyruvate, which we named PyronicSF (Pyruvate Optical Nano-Indicator from CECs Single Fluorophore; DNA sequence is available in Supplementary Fig. S1), was cloned into pcDNA3.1(-) for expression in eukaryotic cells. PyronicSF was targeted to mitochondria or nucleus using pSHOOTER plasmids (ThermoFisher). PyronicSF plasmids are available from Addgene.

### Animals and cultures (mice and flies)

Mixed F1 (C57BL/6J × CBA/J, cultures) were kept in an animal room under Specific Pathogen Free (SPF) conditions at a room temperature of 20 ± 2 °C, in a 12/12 h light/dark cycle with free access to food and water. Procedures were approved by the Centro de Estudios Científicos Animal Care and Use Committee. Mixed cortical cultures (2-3 day-old neonates) were prepared as detailed previously ^46^. Astrocytes in cortical cultures were used at days 8-10. HEK293 cells were acquired from the American Type Culture Collection and cultured at 37 °C in 95% air/5% CO_2_ in DMEM/F12 10% fetal bovine serum. Cultures were transfected at 60% confluence using Lipofectamine 2000 (Gibco) or alternatively, exposed to 5 × 10^6^ PFU of Ad Pyronic (serotype 5, custom made by Vector Biolabs), and studied after 16-24 h. Flies were kept at 25 °C on a standard diet. The fly stock used in this study was apt-Gal4 (P[GMR49G07-GAL4]attP2, Bloomington 45781). The pyruvate sensor expressing fly stocks UAS-PyronicSF and UAS-Mito-Pyronic were generated using the following strategy. The coding sequence of the sensors was amplified by PCR and cloned into a pENTR-vector using pENTR™/D-TOPO^®^ Cloning (ThermoFisher). Then, the sensor coding sequences were cloned via gateway cloning (Gateway LR, Invitrogen) into the vector pUASTattBrfa ^47^, which allows Φintegrase-mediated integration into the fly genome. The resulting vectors have been integrated into the fly genome at landing site attP40.

### Imaging

Cultured cells were imaged at room temperature (22 - 25 °C) in KRH buffer of the following composition (in mM): 136 NaCl, 3 mM KCl, 1.25 CaCl_2_, 1.25 MgSO_4_, 1-2 glucose, 10 HEPES pH 7.4, using an upright Olympus FV1000 confocal microscope equipped with 440, 488 nm and 543 nm laser lines and a 20X water immersion objective (NA 1.0). Pyronic was imaged at 440 nm excitation/480 ± 15 nm (mTFP) and 550 ± 15 (Venus) emissions. mCherry and TMRM were imaged at 543 nm excitation/610 ± 50 emission. Third instar larval *Drosophila* brains were dissected in HL3 buffer (70 mM NaCl, 5 mM KCl, 20 mM MgCl_2_, 10 mM NaHCO_3_, 115 mM sucrose, 5 mM trehalose, 5 mM HEPES; pH 7.1) and placed into a flow-through imaging chamber (Warner Instruments, Hamden, USA). The chamber was fixed onto the stage of a Leica SP8 microscope (Leica Microsystems, Wetzlar, Germany) and brains were superfused with HL3 buffer. Images were acquired using a 40X oil immersion objective (NA 1.4). PyronicSF was imaged at 488 nm excitation/525 ± 25 emission.

### Mathematical modeling

Mitochondrial pyruvate dynamics were simulated using Berkeley Madonna software and the following set of ordinary differential equations:

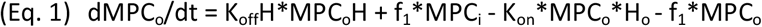

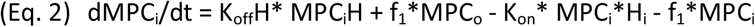

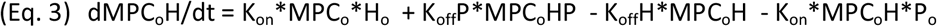

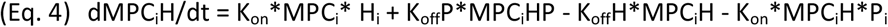

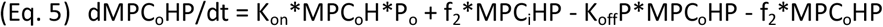

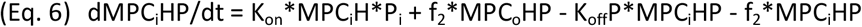

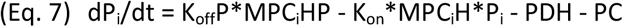

Equations 1 to 6 represent the six conformations of the MPC carrier: outward- and inward-facing, either empty (MPC_o_ and MPC_i_), loaded with a proton (MPC_o_H and MPC_i_H) or loaded with both proton and pyruvate (MPC_o_HP and MPC_i_HP). Equation 7 describes intramitochondrial pyruvate. MPC was set at 3.24 μM; the association constant K_on_ for protons and pyruvate was set at 10^8^ M^−1^*s^−1^; the dissociation constants K_off_H and K_off_P were 20 s^−1^ and 2.12*10^6^ s^−1^; carrier translocation rates f_1_ (empty) and f_2_ (loaded) were set at 200 s^−1^ and 3000 s^−1^. PDH was modeled as a saturable reaction with V_max_ of 1.34 μM* s^−1^ and K_m_ of 10 μM (BRENDA Enzyme Database). Pyruvate dehydrogenase (PDH) was modeled as a saturable reaction with V_max_ of 3.3 μM* s^−1^ and K_m_ of 220 μM (Brenda Enzyme Database). With these parameters and cytosolic and mitochondrial pH values of 7.2 (63 nM) and 7.8 (16 nM ^24^), the simulated carrier displayed accelerated exchange and an apparent zero-trans K_m_ (K_zt_) of 150 μM ^31^. Mitochondrial pyruvate (P_i_) stabilized at 21 μM when cytosolic pyruvate (P_o_) was 33 μM and mitochondrial pyruvate consumption was 1.2 μM* s^−1^ ^26^, with PDH and PC respectively accounting for 76% and 24% of the flux ^27^.

### Statistical Analysis

Statistical analyses were carried out with SigmaPlot software (Jandel). For normally distributed variables, differences were assessed with the Student’s t-test (pairs) and with ANOVA followed by the Tukey-Kramer *ad hoc* test (groups). For variables that failed the normality test, differences were assessed with the Mann-Whitney Rank Sum Test (pairs) or the Kruskal-Wallis one way ANOVA on ranks (groups). *, p < 0.05; ns (non-significant), p > 0.05. The number of experiments and cells is detailed in each figure.

---

**Supplementary Figure S1.**
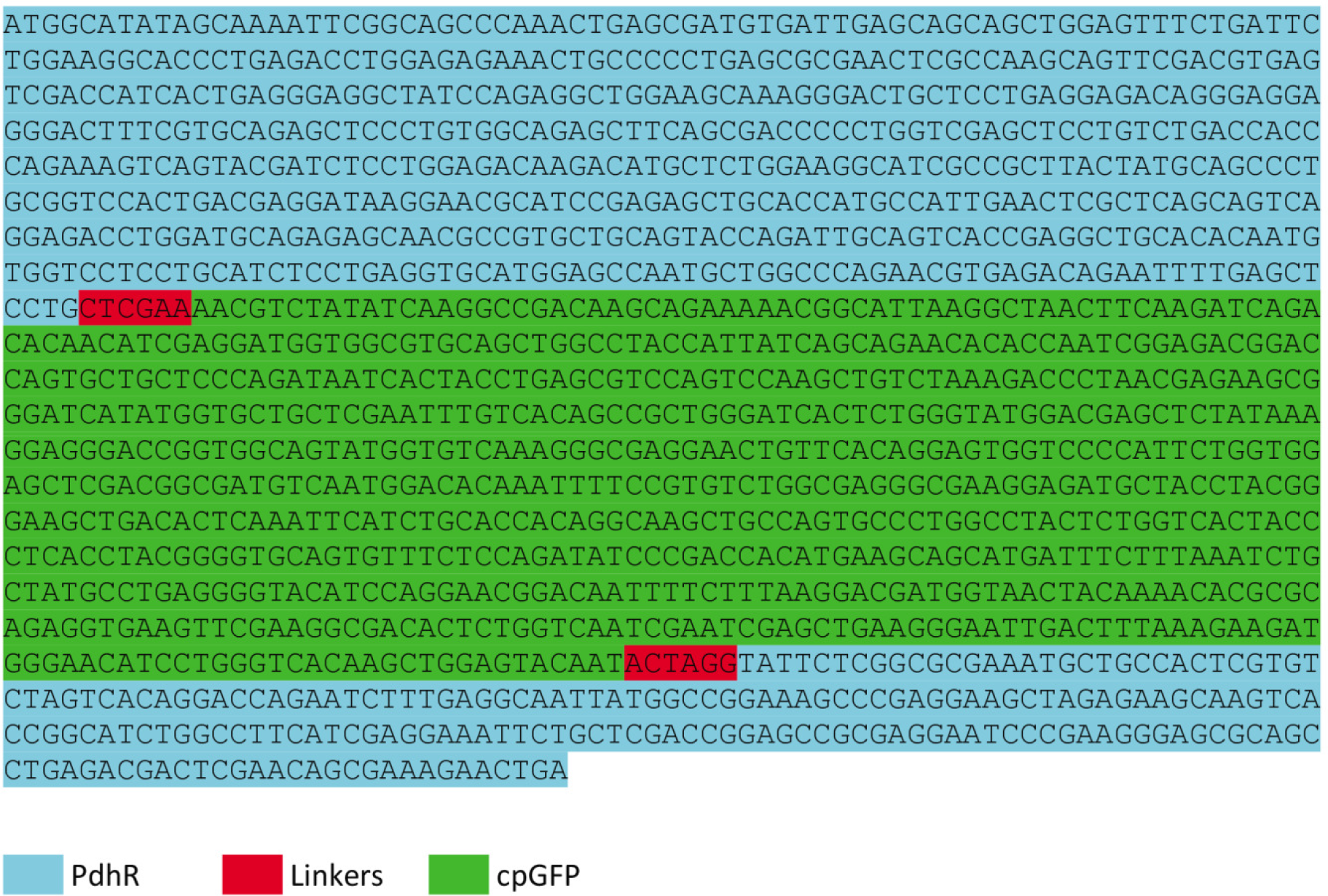
Nucleotide sequence of PyronicSF. DNA sequence of the PyronicSF gene (1500 bp), made from the genes coding for the *Escherichia coli* transcription factor PdhR and the cpGFP, with two linkers as shown. Note that the sequence was codon-optimized for mammalian transcription.

**Supplementary Figure S2.**
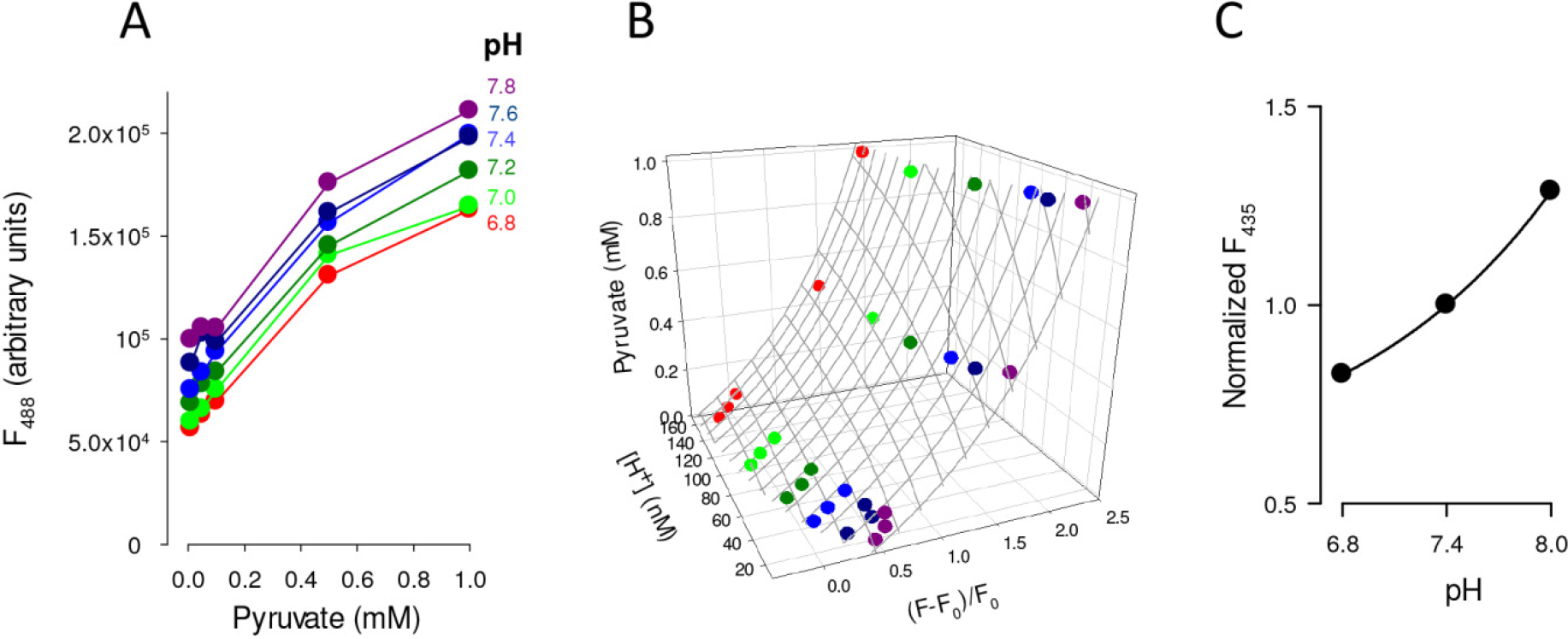
Correction of the effect of pH on PyronicSF. (A) *In vitro* fluorescence of PyronicSF as a function of pyruvate concentration at the indicated pH. Data are means of 3 protein extracts, errors were omitted for clarity. (B) Same data plotted in A, but with fluorescence expressed as F-F_0_/ F_0_ where F_0_ is the fluorescence measured at pH 7.2 in the absence of pyruvate (cytosolic determinations). The gray mesh represents the best fit (r^2^=0.99) of the double hyperbolic function Pyruvate (mM) = 0.942*(F*58.96+F*[H^+^]+1.346*[H^+^]−0.66*58.96-0.66*[H^+^])/(3.367*58.96+3.367*[H^+^]−1.346*[H^+^]+0.66*58.96+0.66*[H^+^]-F*58.96-F*[H^+^]) to the data. At pH 7.8 (mitochondrial determinations) the correcting function is Pyruvate (mM) = 0.942*(F*58.96+F*[H^+^]+0.9032*[H^+^]−0.114*58.96-0.114*[H^+^])/(2.26*58.96+2.26*[H^+^]-0.9032*[H^+^]+0.114*58.96+0.114*[H^+^]-F*58.96-F*[H^+^]). Note that these equations are only valid between 10 μM and 1 mM pyruvate. (C) pH dependency of PyronicSF fluorescence measured at 435 nm excitation, at which the sensor is insensitive to pyruvate (Fig. 1B). The line shows the best fit of the equation F_435_ = 0.55 + 0.001*e^(0.82*pH) to the data. This sensitivity may be used to control for possible pH changes and correct them using the equation in B, as an alternative to parallel measurements with a pH probe.

**Supplementary Figure S3.**
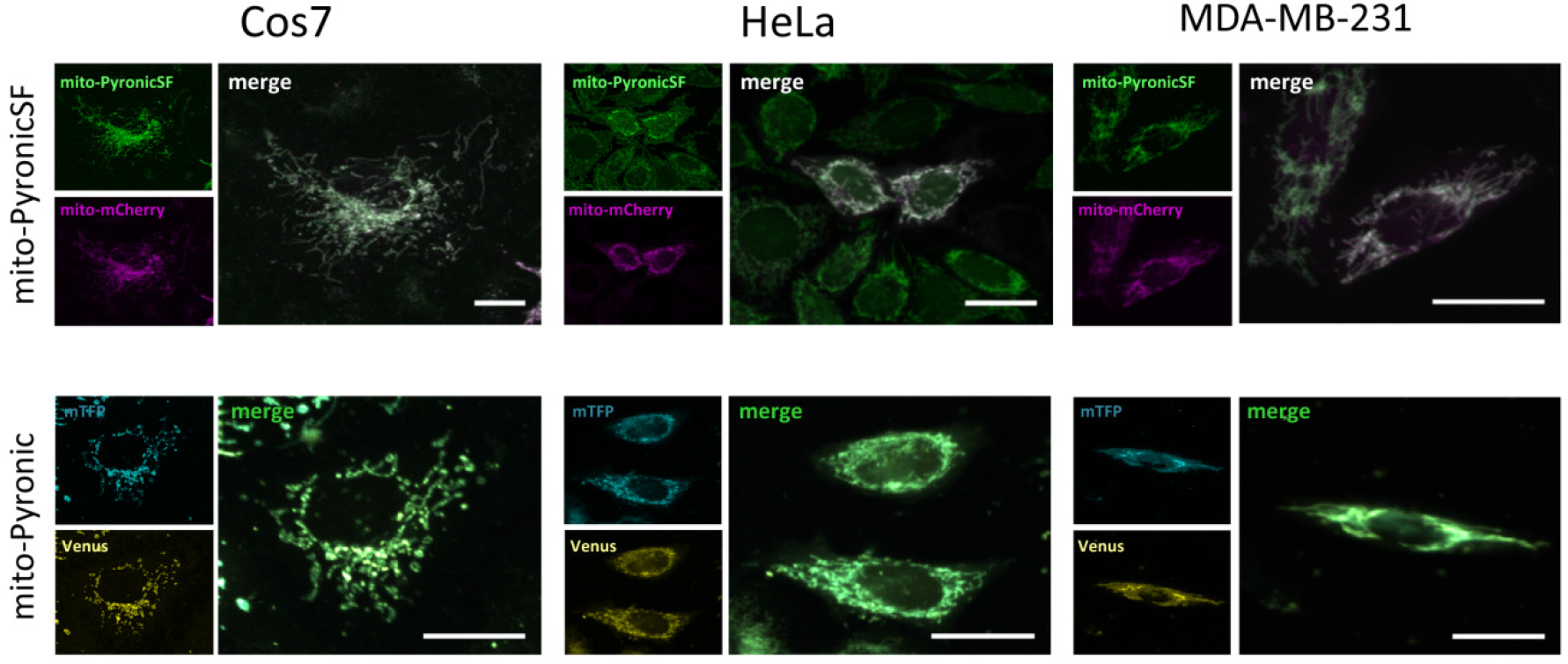
Expression of mito-PyronicSF and mito-Pyronic in various cell types. Mito-PyronicSF (top row) and mito-Pyronic (bottom row) were expressed in Cos7, HeLa and MDA-MB-231 cells. Mito-PyronicSF is shown in green and mitochondrial mCherry is shown in magenta. The components of the FRET sensor Pyronic are shown in cyan (mTFP) and yellow (Venus). Scale bars represent 20 μm.

**Supplementary Figure S4.**
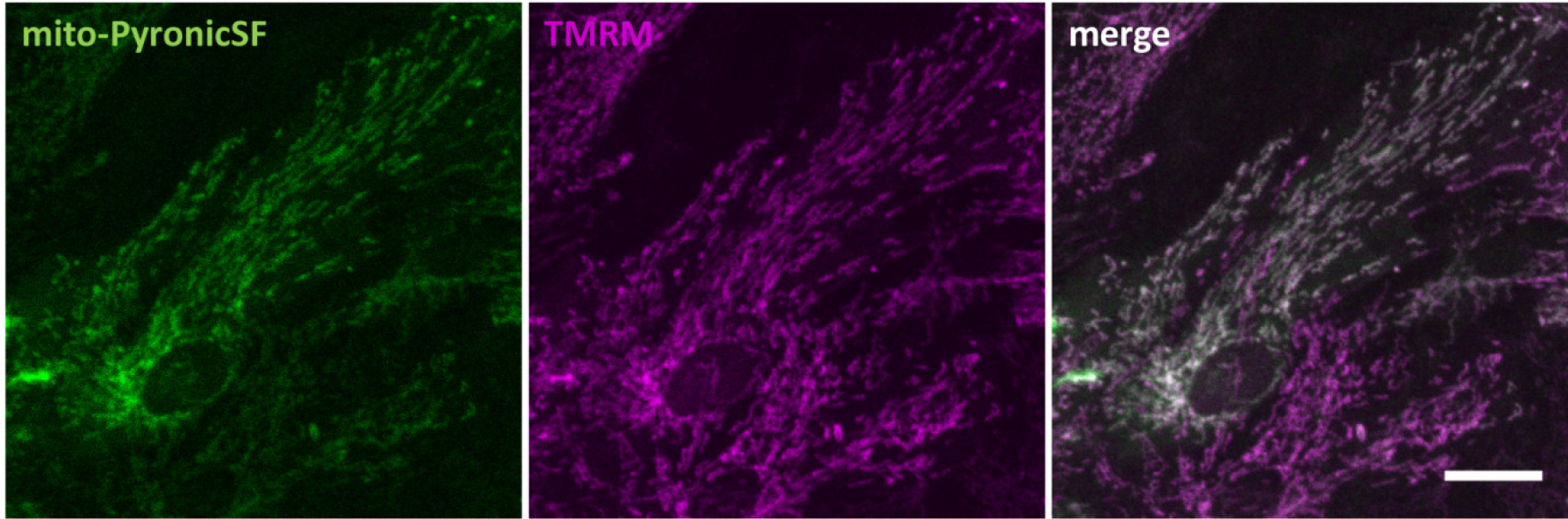
Mitochondrial localization of mito-PyronicSF in astrocytes. A culture of astrocytes expressing mito-PyronicSF was loaded with 400 nM TMRM for 30 s. Images of mito-Pyronic (green) and TMRM (magenta) are shown, as well as their merged image. Scale bar represents 20 μM.

**Supplementary Figure S5.**
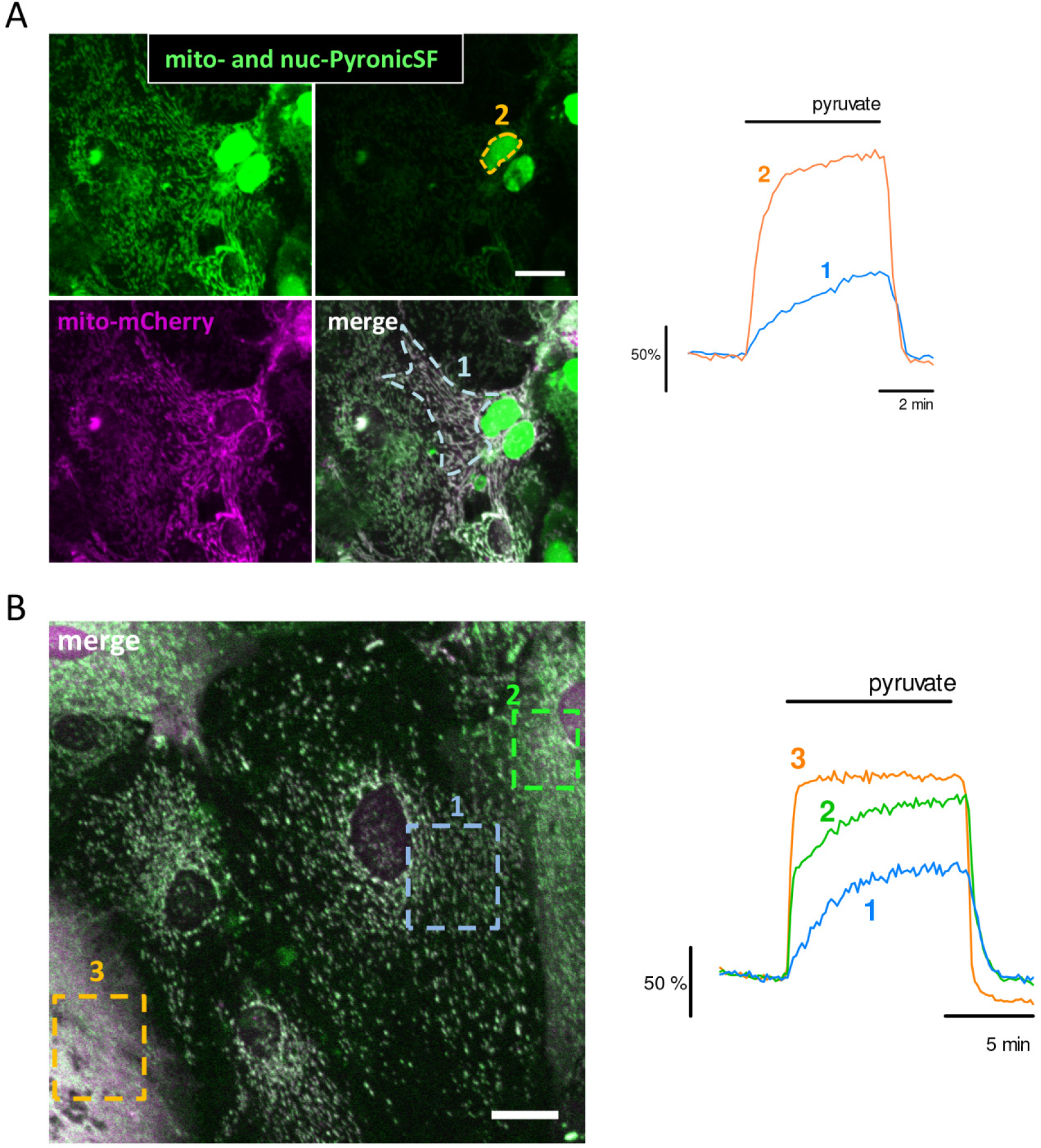
Mitochondrial pyruvate uptake is slower than cytosolic pyruvate uptake. (A) Astrocytes co-expressing mito-PyronicSF and mito-mCherry by means of adenoviral transduction were transfected with nuc-PyronicSF. To avoid photomultiplier saturation, filtered images were used for quantification of the nuclear sensor (top right). Scale bar represents 10 ⍰m. The graph shows the response to 10 mM pyruvate of two regions of interest drawn on the same cell, corresponding to mitochondria (#1, blue) and nucleus (#2, orange). (B) Astrocytes co-expressing mito-PyronicSF and mito- mCherry. Regions of interest were drawn on three cells showing variable levels of diffuse cytoplasmic fluorescence (“leak”), ranging from undetectable (#1, blue), to intermediate (#2, green) and maximum (#3, orange). Scale bar represents 10 μm. The responses to the three regions of interest to 10 mM pyruvate are shown in the graph.

**Supplementary Figure S6.**
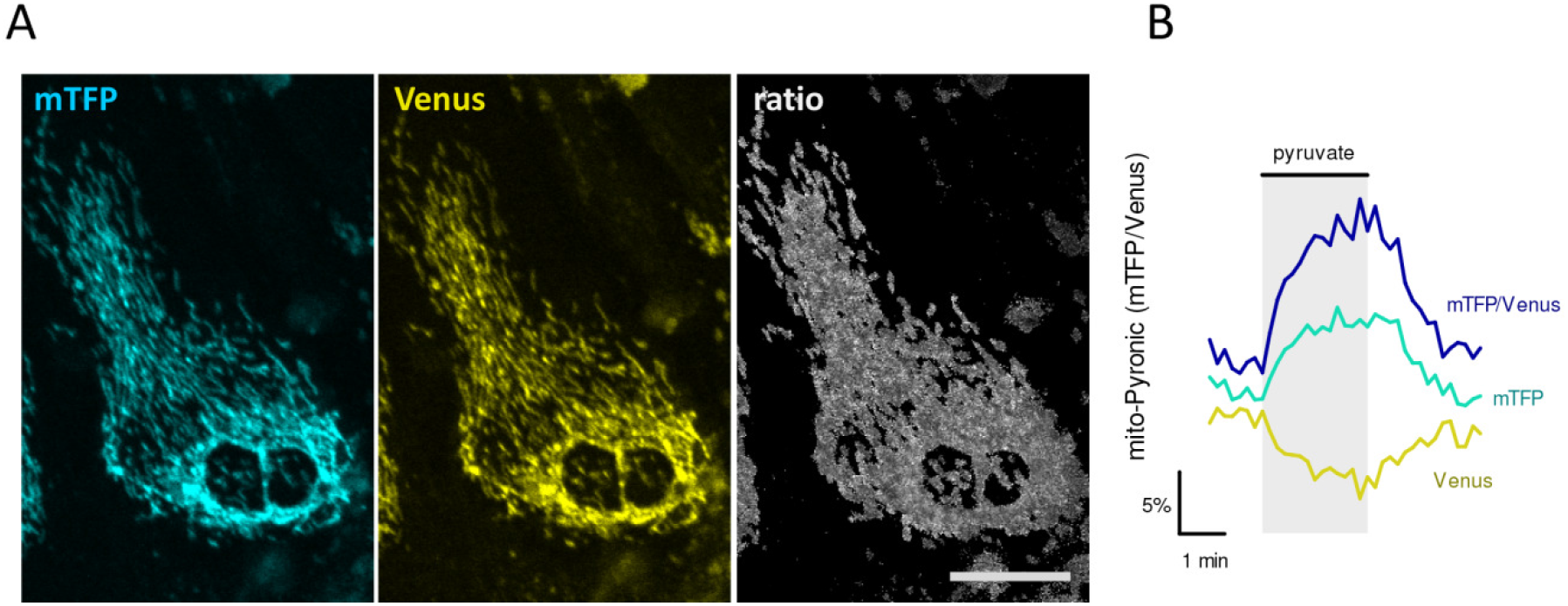
Functional expression of the FRET sensor Pyronic in mitochondria. (A) Expression of Pyronic in astrocytic mitochondria was achieved with four copies of the destination sequence of cytochrome oxidase. Images show mTFP, Venus and the ratio between mTFP and Venus. Bar represents 10 μm. (B) Response of the astrocytes shown in A to 10 mM pyruvate.

